# Single molecule tracking of AMPA receptors shows the role of synaptic insertion during maintenance of chemical LTP

**DOI:** 10.1101/2021.02.04.429665

**Authors:** Chaoyi Jin, Sung-Soo Jang, Pinghua Ge, Hee Jung Chung, Paul Selvin

## Abstract

Long term potentiation (LTP) likely contributes to memory formation. Early expression of LTP involves insertion of AMPA receptors (AMPARs) to the extrasynaptic membrane followed by their lateral diffusion into the synaptic membrane. However, whether a similar mechanism mediates the maintenance of LTP is unclear. Using single-molecule microscopy, we quantified that 6 GluA1- and 11 GluA2-containing endogenous AMPARs were added per synapse in cultured hippocampal neurons at 20 min following chemical LTP (cLTP) induction for 10 min, resulting in a 54% increase for both subunits. Single molecular tracking of transfected subunits revealed that the number of exocytosed subunits at the synapse increased by 15-18% from 5 to 20 min following cLTP induction, but their lateral exchange between synaptic and extrasynaptic membranes was minimal. These findings suggest that cLTP maintenance is contributed largely by synaptic insertion of AMPARs rather than the surface diffusion of exocytosed AMPARs from extrasynaptic to synaptic regions.

---

AMPA receptors (AMPARs) are ionotropic glutamate receptors that mediate fast excitatory synaptic transmission in the central nervous system (CNS) (*1*–*3*). An AMPAR has four subunits (GluA1-A4) that are predominantly located at the post synaptic density (PSD) (*2, 4*). Long term potentiation (LTP) of excitatory synaptic strength is critical for learning and memory, whereas its disruption is associated with memory impairment. It is therefore important to understand the molecular mechanism underlying how LTP is expressed at the beginning (early expression at 0-10 min after stimulation) and maintained for longer term (maintenance at >10 min) (*5*).

Extensive studies have demonstrated that an increase in the conductance and number of postsynaptic AMPARs mediates the expression of LTP (*3, 5*–*10*). It is widely believed that AMPARs exocytose into the extrasynaptic plasma membrane from intracellular stores and move into the synapse by surface diffusion (*3*), although direct insertion into the synaptic region has also been proposed (*3, 5, 6, 11*). A recent study has shown that early expression of hippocampal LTP involves lateral diffusion of surface AMPARs into the synapse (*12*). However, whether this same mechanism is also required for maintaining LTP expression, remains elusive (*7, 11, 13*).

This knowledge gap is in part due to the challenges in detecting the precise insertion site, the number, and the trafficking of AMPARs in the synaptic and extrasynaptic plasma membrane during LTP expression in a living neuron. Tagging with fluorescent proteins fails because of their poor photostability (*7*). Labeling with commercial quantum dots (QDs) overcomes poor photostability but is not ideal because of their large size (>20 nm) (*4, 14*–*16*) that could alter receptor mobility. Similarly, the use of both primary and secondary antibodies is problematic due to their large size (*4, 13*).

Here, we observed surface AMPARs on a synapse-by-synapse basis during the maintenance of chemical LTP (cLTP) with single-molecule sensitivity by using primary antibodies conjugated with stable fluorophores or with streptavidin-conjugated small QDs. In our study, endogenous and transfected AMPARs are counted at excitatory synapses in cultured hippocampal neurons before and after inducing cLTP and the surface diffusion of *newly inserted* AMPARs was tracked during the maintenance phase of cLTP. We found that AMPARs were exocytosed into both synaptic and extrasynaptic membrane of cultured hippocampal neurons upon cLTP induction; however, we also found that the exocytosed AMPARs accumulate at synapses during the maintenance period of LTP without affecting their net flow from the extrasynaptic to synaptic membrane.

## Results

To induce cLTP, we treated dissociated culture of hippocampal neurons with NMDA receptor (NMDAR) co-agonist glycine (200 µM) in Mg^2+^-free solution. This well-established “cLTP induction” protocol selectively activates synaptic NMDARs and rapidly increases synaptic AMPAR expression (*17, 18*). Whole-cell patch-clamp recording of hippocampal pyramidal neurons revealed that the amplitude of mEPSCs, indicative of postsynaptic response, was significantly increased at 20 min after application of glycine (23.2 pA ± 2.7) compared to vehicle control (17.1 ± 1.4; p < 0.05), confirming the expression of cLTP (*17*) (Fig. S1).

To observe *endogenous* AMPARs on the same synapses before and after cLTP induction, the neurons were transfected with a PSD protein, Homer1c, as a reference for the excitatory synapse. Homer1c was coupled to mGeos, a photoactivatable fluorescent protein (emission maximum at 515 nm), that can be localized with super-resolution microscopy to ∼ 20 nm (*19*). The synaptic region was defined as being 0∼0.5 µm from the center of Homer1c, whereas the extrasynaptic region was defined as being >0.5 µm from the center of Homer1c (*4*). Before cLTP induction, GluA1 or GluA2 were first saturated with Cy3b-conjugated (emission ∼ 572 nm) primary antibodies (“pre-cLTP”, Fig. 1A-B). At 20 min after cLTP induction for 10 min, exocytosed GluA1 or GluA2 were then labeled with CF633-conjugated (emission = 650 nm) primary antibody (“post-cLTP”, Fig. 1A-B). Finite kinetics of antibody-binding prevented neurons from being imaged earlier than this after cLTP induction.

**Figure 1:**
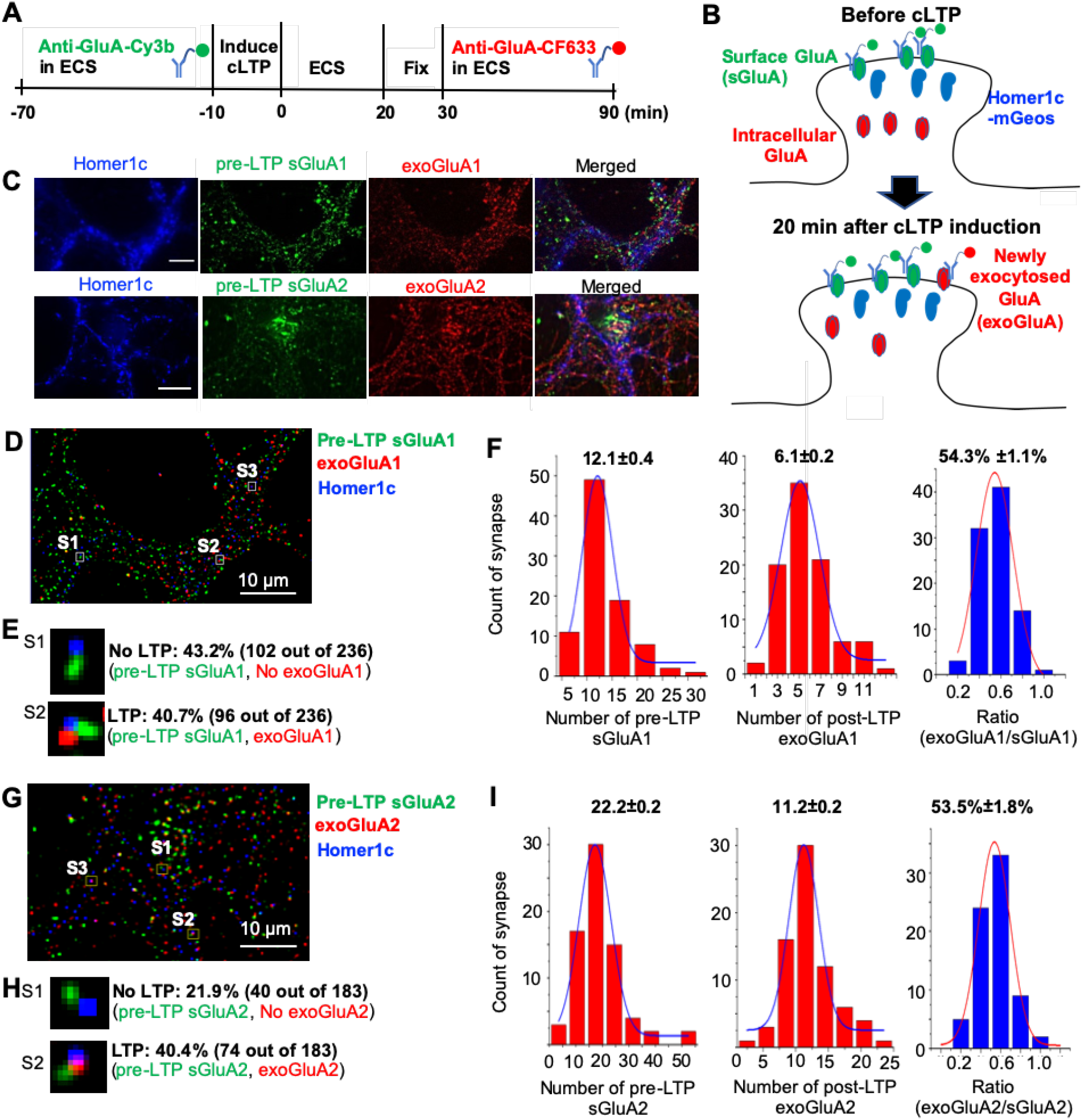
The induction of cLTP increases synaptic insertion of *endogenous* AMPAR subunits GluA1 and GluA2 in cultured primary hippocampal neurons. **(A)** Schematic of an experimental procedure to induce cLTP and label surface AMPARs in cultured primary hippocampal neurons transfected with a PSD marker Homer1c-mGeos. Endogenous GluA1 or GluA2 on neuronal plasma membrane before cLTP induction (sGluA1, sGluA2) were saturated with anti-GluA1-Cy3b or anti-GluA2-Cy3b for 60 min in extracellular control solution (ECS). The cLTP was induced by incubating neurons for 10 min in glycine-induced synaptic potentiation (GISP) solution containing 200 μM glycine in ECS without Mg^2+^. After 20 min in ECS, neurons were fixed and exocytosed GluA1 or GluA2 were labeled with 60 nM anti-GluA1-CF633 or anti-GluA2-CF633, respectively, for 60 min in ECS. **(B)** Schematic of endogenous AMPAR subunit labeling outcome. **(C)** Representative images of dendrites showing transfected Homer1c-mGeos (blue), surface GluA1or GluA2 prior to cLTP induction (green), and the newly exocytosed GluA1 or GluA2 after cLTP induction (red). Scale Bar: 10 μm. **(D-F)** Analysis GluA1. **(D)** A merged image of GluA1 from Fig. 1C containing 3 different synapses (Synapses 1-3). **(E)** Zoomed images of Synapse 1 and Synapse 2 from Fig. 1D. Zoomed image of Synapse 3 is shown in Fig. S2A. **(F)** Quantification of synapses that show pre-existing surface GluA1 prior to cLTP and newly inserted GluA1 after cLTP induction, in relation to the number of exocytosed GluA1. **(G-I)** Analysis GluA2. **(G)** A merged image of GluA2 from Fig. 1C containing 3 different synapses (Synapses 1-3). **(H)** Zoomed images of Synapse 1 and Synapse 2 from Fig. 1G. Zoomed image of Synapse 3 is shown in Fig. S2A. **(I)** Quantification of synapses that show pre-existing surface GluA2 prior to cLTP induction and newly inserted GluA2 after cLTP induction, in relation to the number of exocytosed GluA2. A total of 236 synapses for GluA1 was measured from 2 neurons in 2 independent experiments. A total of 183 synapses for GluA2 was measured from 2 neurons in 2 independent experiments.

We observed 12.1 ± 0.4 endogenous surface GluA1 and 22.2 ± 0.2 endogenous surface GluA2 in total (extrasynaptic and synaptic) under “pre-LTP” condition (sGluA’s, Fig. 1C-I). Since 70% of total GluA2 proteins is reported to be synaptic (*4*), we calculated 22*0.7 = 15.4 as the number of synaptic GluA2. Since the majority of AMPARs in hippocampal neurons are GluA1/A2 and GluA2/A3 heteromers, our measurement of GluA2 number is consistent with 15 synaptic AMPARs observed in fixed brain tissue by electron microscopy (*14*). This consistency validated our fluorescence measurements.

At 20 min post-cLTP induction, heterogeneity was detected. For GluA1 and GluA2, 43.2% and 21.9% of synapses did not respond to glycine treatment, respectively (Fig. 1E, synapse S1). These synapses had surface GluA1 and GluA2 before cLTP but lack newly exocytosed subunits after cLTP induction. Among the synapses that responded to glycine treatment (56.8% for GluA1 and 88.1% for GluA2), 40.7% had surface GluA1 and 40.4% had surface GluA2 before cLTP and acquired newly exocytosed GluA1 after cLTP induction (Fig. 1E, synapse S2), whereas 16.1% and 37.7% lacked surface GluA1 and GluA2, respectively, before cLTP but gained newly exocytosed GluA1 after cLTP induction (Fig. 1E, S2, synapse S3).

In synapses that already contained GluA1 and GluA2 at pre-LTP, cLTP induction increased the number of endogenous surface GluA1 by 54.3% from 12.1 ± 0.4 to 18.2 ± 0.6 (Fig. 1F) and surface GluA2 by 53.5% from 22.2 ± 0.2 to 33.4 ± 0.4 (Fig. 1I). The extent of increase in the number of GluA1 and GluA2 on these synapses (6.1 ± 0.2 and 11.2 ± 0.2, respectively, Fig. F, I) were similar to that in the synapses that lacked surface AMPAR subunits before cLTP (6.1 ± 0.2 and 11.7 ± 0.7, respectively, Fig. S2). This suggests that the increase in the number of AMPARs on the synaptic plasma membrane upon cLTP induction is dependent on AMPAR subunit, but not on the type of synapses, i.e. whether it had sGluA’s before cLTP, or not (Fig. 1, S2). The greater increase in GluA2 than GluA1 may likely be explained by the fact that GluA1 antibody only detects GluA1/A2 receptors, while GluA2 antibody can detect both GluA1/A2 and GluA2/A3 receptors (*17*).

Although the increase in synaptic AMPARs occurs during early expression and maintenance of LTP (*17, 20*), it is unclear whether they are inserted into the synaptic or extrasynaptic regions during cLTP maintenance (*11*) the precise number of exocytosed AMPARs during cLTP maintenance has not been determined (*3, 11, 17, 20*). To investigate this, we transfected neurons with Homer1-mGeos and GluA1 or GluA2 containing an acceptor peptide-tag at the extracellular domain (GluA-AP) which was biotinylated by co-transfected biotin ligase (BirA) (*21*). We first incubated transfected neurons with unlabeled streptavidin (SA) to saturate biotinylated GluA1 or GluA2 on the surface before cLTP induction (Fig. 2A-B, Fig. S3). After cLTP induction, newly inserted biotinylated GluA1 or GluA2 was labeled with SA-conjugated Atto647N for 5 min and imaged immediately at 5 min post-cLTP induction (Fig. 2A-C, Fig. S3-4), or imaged 15 min later at 20 min post-cLTP induction (Fig. 2A-B, D). However, high concentration of SA (50 nM of free SA or 500 nM of SA-dye conjugates) has been shown to crosslink surface AMPARs and other proteins (*12, 22*); therefore, we used much lower concentration of SA-Atto647N (0.6 nM) (Fig. 2, 3), which we have previously verified not to induce cross-linking of AMPARs (*4*).

**Figure 2.**
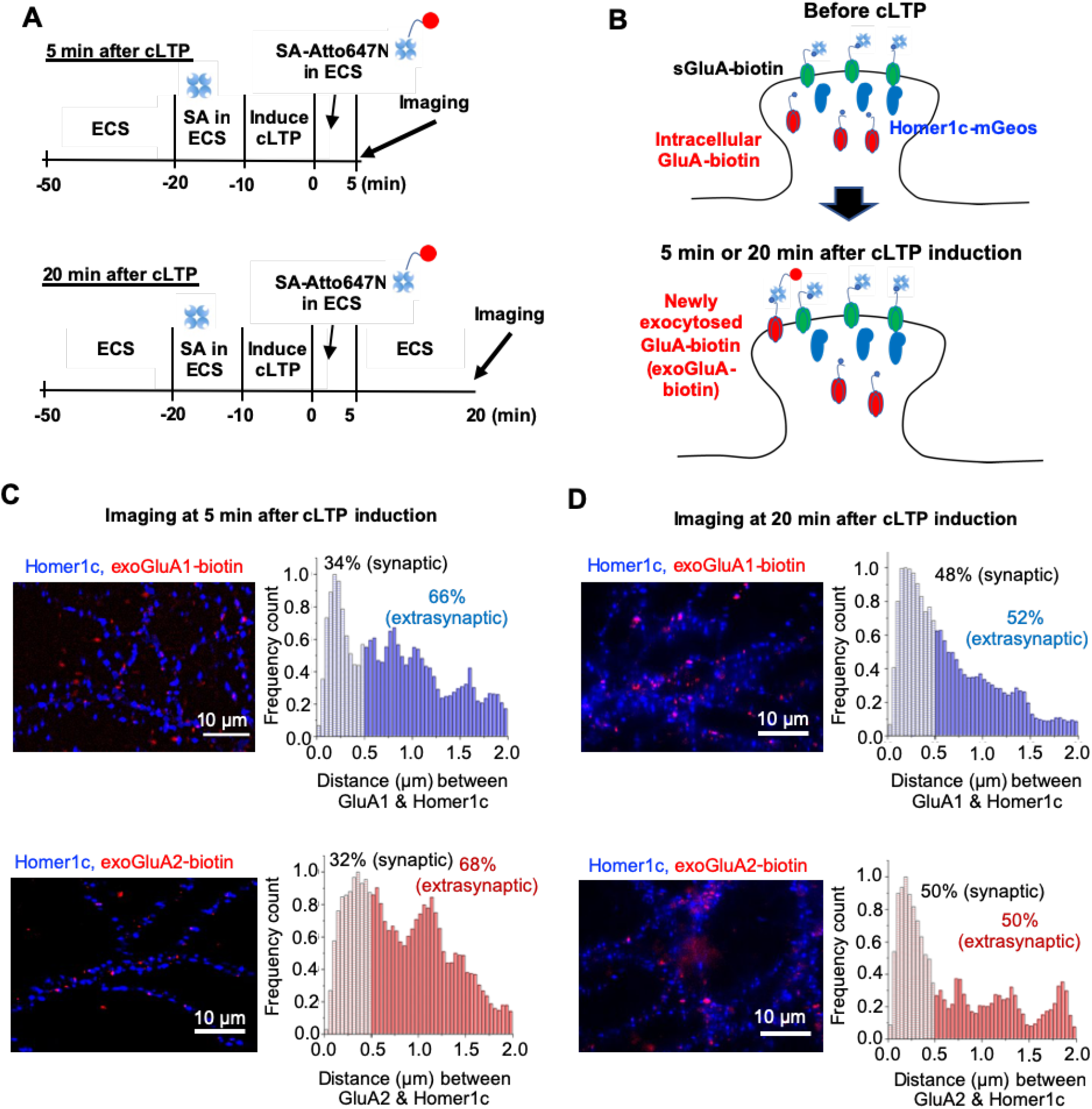
The induction of cLTP increases exocytosis of transfected AMPAR subunits and the percentage of exocytosed subunits at excitatory synapses increases from 5 to 20 min post cLTP induction. **(A)** Schematic of an experimental procedure to label transfected AMPARs that are newly inserted to the plasma membrane at 5 and 20 min after cLTP induction. Primary hippocampal cultured neurons were transfected with a PSD marker Homer1c-mGeos and GluA1 or GluA2 containing an acceptor peptide tag at the extracellular domain which were biotinylated by co-transfected biotin ligase (GluA-biotin). Neurons were incubated with 10 nM free streptavidin (SA) for 10 min to saturate pre-existing surface GluA1-biotin and GluA2-biotin before cLTP induction. After treating neurons in GISP solution for 10 min to induce cLTP, 0.6 nM SA-Atto647N were applied to neurons for 5 min to label newly inserted GluA-biotin and imaged immediately (top procedure) or 15 min later at 20 min after cLTP (bottom procedure). **(B)** Schematic of labeling outcome of biotinylated AMPAR subunit that are newly inserted upon cLTP induction. The receptor is defined to be synaptic if this distance is less than 0.5 um; the receptor is extrasynaptic if this distance is 0.5-2 um. **(C)** (Left) Representative images of dendrites showing transfected Homer1c-mGeos (blue) and the newly exocytosed GluA1 after cLTP induction (red, exoGluA1). (Right) Frequency count histogram on *transfected* GluA1 in relation to the distance between the center of Homer1c (a PSD marker) and newly inserted GluA1 at cLTP at 5 min and 20 min post cLTP induction. A total of 396 synapses for GluA1 was measured from 5 neurons at 5 min post cLTP induction (2 independent experiments). A total of 565 synapses for GluA1 was measured from 5 neurons at 5 min post cLTP induction (3 independent experiments). **(D)** (Left) Representative images of dendrites showing transfected Homer1c-mGeos (blue) and the newly exocytosed GluA2 after cLTP induction (red, exoGluA2). (Right) Frequency count histogram on *transfected* GluA2 in relation to the distance between the center of Homer1c (a PSD marker) and newly inserted GluA2 at cLTP at 5 min and 20 min post cLTP induction. A total of 1012 synapses for GluA2 was measured from 9 neurons at 20 min post cLTP induction (3 independent experiments). A total of 589 synapses for GluA2 was measured from 6 neurons at 20 min post cLTP induction (2 independent experiments). The synaptic percentage of GluA1 and GluA2 increases 14% and 18%, respectively, when imaged at 20 min post cLTP induction compared to 5 min post cLTP induction.

**Figure 3.**
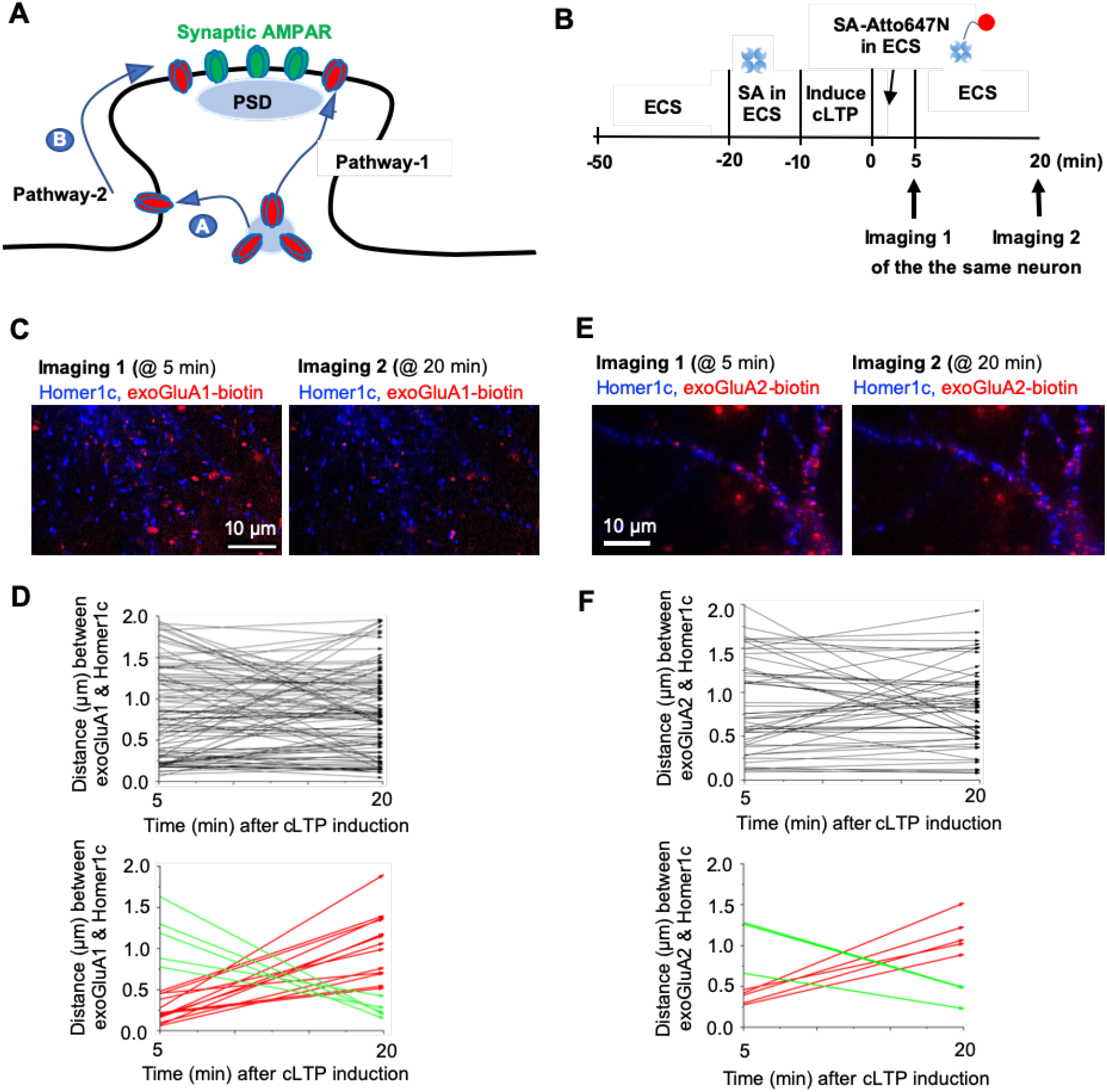
Lateral movement of newly inserted AMPARs from extrasynaptic region to synaptic regions is minimal during from 5 min to 20 min post cLTP induction. **(A)** Hypothesis by which synaptic addition of AMPARs can occur for LTP maintenance. Pathway-1 involves direct insertion of AMPAR into a synaptic region before entering the PSD via membrane diffusion over a short distance. Pathway-2 involves exocytosis of AMPARs to the extrasynaptic region (route A) followed by lateral surface diffusion to the PSD (route B). **(B)** Schematic of an experimental procedure to label transfected AMPARs that are newly inserted to the plasma membrane and track their lateral diffusion from 5 min to 20 min after cLTP induction. Neurons were incubated with 10 nM free streptavidin (SA) for 10 min to saturate preexisting surface GluA1-biotin and GluA2-biotin before cLTP induction. After treating neurons in GISP solution for 10 min to induce cLTP, 0.6 nM SA-Atto647N were applied to neurons for 5 min to label newly inserted GluA-biotin and imaged immediately (Imaging 1). After 15 min (at 20 min after cLTP induction), the same neurons were imaged again to track the movement of each individual GluA1 or GluA2 (Imaging 2). **(C, E)** Representative images of dendrites showing transfected Homer1c-mGeos (blue) and the newly exocytosed GluA1 **(C)** and GluA2 **(E)** after cLTP induction (red, exoGluA1 or exoGluA2) at 5 min after cLTP induction (Imaging 1) and 20 min after cLTP induction (imaging 2). **(D, F)** Distance between exocytosed GluA1 **(D)** and GluA2 **(F)** and the center of Homer1c at 5 and 20 min after cLTP induction at the synapse that underwent cLTP indicated by the increase in the volume of Homer1c-mGeos intensity from 5 min to 20 min (See Fig. S8). The end of an arrow stands for the moving direction of the subunit. The top graphs show all traces (black). The bottom graphs display the traces extracted from the top graphs, which show the subunits moving from synaptic to extrasynaptic region (red), or those from extrasynaptic to synaptic region (green). A total of 100 synapses for GluA1 was measured from 3 neurons (2 independent experiments). A total of 53 synapses for GluA2 was measured from 2 neurons (2 independent experiments).

We measured the distances between transfected subunit and the center of Homer1c, and defined AMPAR as synaptic when it is located ≤0.5 μm from the center of Homer1c-mGeos (*4*). We found that exocytosed GluA1 and GluA2 in both synaptic and extrasynaptic membrane (Fig. 2C-D). At 5 min post-cLTP induction, 34% of exocytosed GluA1 and 32% of exocytosed GluA2 subunits were synaptic (Fig. 2C). At 20 min post-cLTP induction, the exocytosed GluA1 and GluA2 were 48% and 50% synaptic, respectfully (Fig. 2D). This percentage increase in synaptic AMPARs corresponds to the increase in the *number* of synaptic AMPARs (Fig. S5, Table S1-2). During our experimental duration, AMPARs exocytosed upon cLTP induction accounted for >95% of all the synapses (Fig. S3-4), making their basal exocytosis negligible.

We hypothesize that the increase in the number of synaptic AMPARs during cLTP maintenance could be achieved by two pathways (Fig. 3A). Pathway-1 is the AMPAR exocytosis at or near the PSD (Fig. 3A); Pathway-2 is the insertion of AMPARs at extrasynaptic domain followed by their lateral movement along the plasma membrane to the synaptic site (Fig. 3A, routes A and B). To test if Pathway-2 contributes to the observed increase in synaptic AMPARs during cLTP maintenance, we labeled only exocytosed GluA1 or GluA2 post-cLTP induction with SA-Atto647N for 5 min as in Fig. 2 (Fig. 3B) and tracked the location of the *same* subunit at 5 and 20 min post-cLTP induction at the synapse in which cLTP was expressed (Fig. 3B-3D). The synapses undergoing cLTP was identified by the increase in Homer1c fluorescent intensity (Fig. S6), since LTP is reported to associate with a gradual enhancement in synaptic content of Homer1c (*23*).

We find that the exchange rate between the synaptic (≤ 0.5 μm from the center of Homer1c) and extrasynaptic regions (0.5-2 μm from the center of Homer1c) remains unchanged for both the synaptic and extrasynaptic population over the course of 5 to 20 min post-cLTP induction. The same conclusion was obtained when the synaptic region was defined as being ≤0.6 μm from the center of Homer1c (Fig. S7), ensuring that the result did not depend on the exact definition of the distance between synaptic and extra-synaptic regions. We find that out of 100 GluA1, there were 13 GluA1 moving from synaptic to extrasynaptic region (red arrows), and 5 moving from extrasynaptic to synaptic region (green arrows) (Fig. 3C-D). Similarly, out of 53 GluA2, there were 5 GluA2 moving from synaptic to extrasynaptic region (red arrows), and 3 GluA2 moving from extrasynaptic to synaptic region (green arrows) (Fig. 3E-F). Furthermore, we found that the extrasynaptic GluA1 and GluA2 subunits were ∼ 1.5 and ∼ 2.2 more diffusive at 5 min and 20 min post cLTP induction than the synaptic ones (Table S3), consistent with the notion that synaptic AMPARs are tethered to PSD, whereas extrasynaptic AMPARs are not bound to PSD proteins and thus are more mobile (*4*).

To validate our results with SA-Atto647N, we used small QDs conjugated with monomeric SA (mSA-sQDs), which show ∼85% probability of labeling AMPARs within excitatory synapses, due to its compact size (∼12 nm). The sQDs were imaged at super-resolution with ∼10-20 nm accuracy (*16*) and their photostability allows the tracking of the same subunits every 5 min from 5-20 min post-cLTP induction *(4)* (Fig. S8). We focused on GluA2 because most AMPARs in the hippocampal neurons contain GluA2 (*6*). Out of 24 GluA2 exocytosed at 5 min post cLTP induction, there were 3 GluA2 moving from synaptic to extrasynaptic region, and 2 moving from extrasynaptic to synaptic region (Fig. S8). The net change was 1 GluA2 moving from synaptic to extrasynaptic region during 5-20 min post-cLTP induction. New patterns of motion were rare since only 2 out of 24 GluA2 “hopped” between the two regions (Fig. S8). These results were consistent with our data using SA-Atto647N (Fig. 3) and suggests that the surface diffusion of exocytosed AMPARs is minimal between the synaptic and extrasynaptic regions during cLTP maintenance.

## Discussion

In this study, we present quantitative measurement of the numbers, percentages, and movements of synaptic GluA1 and GluA2 during cLTP maintenance in living hippocampal cultured neurons at the single-synapse and single-molecule level. Here we performed super-resolution microscopy with smaller sQDs (<12 nm) or organic fluorophores (∼ 5 nm) to track the number of newly inserted AMPARs and their distribution before and after cLTP induction. Using these small probes, we have recently demonstrated that AMPARs labeled with such small probes are mostly synaptic under basal conditions (*4*) whereas AMPARs labeled with large commercial QDs (>20 nm in diameter) (*24*) are not (*4, 16*).

In this study, we discovered that the number of the newly exocytosed AMPARs at 20 min post-cLTP induction is dependent on whether or not the subunit had GluA’s before cLTP, i.e. subunit identity (Fig. 1). At 5 min after cLTP induction treatment, the newly exocytosed AMPARs are found in both synaptic and extrasynaptic membranes (Fig. 2). Extrasynaptic exocytosis of AMPARs upon stimulation of neuronal activity agrees with previous publications (*11, 20, 25, 26*). However, these reports did not specify what proportion of the AMPARs were inserted extrasynaptically during LTP maintenance (*11, 20, 25, 26*). We found that ∼66% of the newly exocytosed AMPARs is in the extrasynaptic region, and the remaining ∼33% is in the synaptic region at 5 min after cLTP induction. At 20 min following cLTP, 50% of the total AMPARs were localized at the synaptic region. This suggests that synaptic AMPARs increases during the cLTP maintenance.

Multiple studies used a combination of electrophysiology and imaging of AMPARs tagged with a pH-sensitive fluorescent protein or labeled with large commercial QDs, show that lateral diffusion of surface AMPARs from the extrasynaptic site to the synaptic region occurs during early expression of LTP (*2, 3, 12, 24, 27, 28*). These studies have led to the prevailing ‘slots’ hypothesis (*3, 24*). Through the results of big quantum dots, this has been extended to argue that AMPARs are majorly extrasynaptic and highly mobile under basal conditions but get captured by the slot proteins at synapse (*24*). The crosslinking of surface AMPARs has recently been shown to significantly decrease *early* expression of LTP (*12*), further demonstrating that the lateral movement of pre-existing surface AMPARs to the potentiated synapse mediates early expression of LTP. However, crosslinking does not eliminate the sustained expression of LTP (*12*). Rather, this was blocked by Tetanus Toxin light chain which inhibits postsynaptic membrane fusion events (*12*), suggesting that AMPAR exocytosis is important for the sustained expression, i.e., maintenance, of LTP.

Consistent with this notion, our labeling and tracking of newly exocytosed AMPARs using 2 different probes (SA-Atto647N and mSA-sQD) revealed that the net exchange of AMPARs between synaptic and extrasynaptic regions is very small (<4% of the exocytosed AMPAR after cLTP), despite a larger increase (15-18%) in synaptic AMPARs during cLTP maintenance from 5 to 20 min after cLTP induction (Fig. 2-3, S8). These results suggest that the surface diffusion of newly inserted AMPARs from extrasynaptic to synaptic regions is not the main cause for the increase in synaptic AMPARs during cLTP maintenance in dissociated primary neuronal cultures.

We propose that either postsynaptic insertion of AMPARs and/or an increase in their packing density at the synaptic region may mediate cLTP maintenance. The exocytosis of endosomes containing AMPARs and transferrin receptors occurs immediately adjacent to the PSD within dendritic spines (*26*), yet only AMPARs bind to synaptic anchoring proteins and are retained at excitatory synapses (*26*). Importantly, a nanoscale intrasynaptic movement of AMPARs induces their enrichment at the dense nanodomains of PSD-95: this has been observed following LTP and can potentiate synaptic currents (*29, 30*). Therefore, it is also possible that exocytosed AMPARs become further concentrated at excitatory synapses by moving a nanoscale distance and binding to synaptic anchoring proteins during cLTP maintenance (27). In summary, our quantitative single molecule tracking of AMPARs with smaller and photostable probes has provided a mechanistic insight on how cLTP is maintained. Exploring these novel mechanisms in the LTP maintenance of intact hippocampal circuitry warrants for future research.

## Acknowledgments

This work was supported in part by NIH grants NS100019, NS097610 and NSF PHY 1430124.

## Supplementary Figures

**Figure S1.**
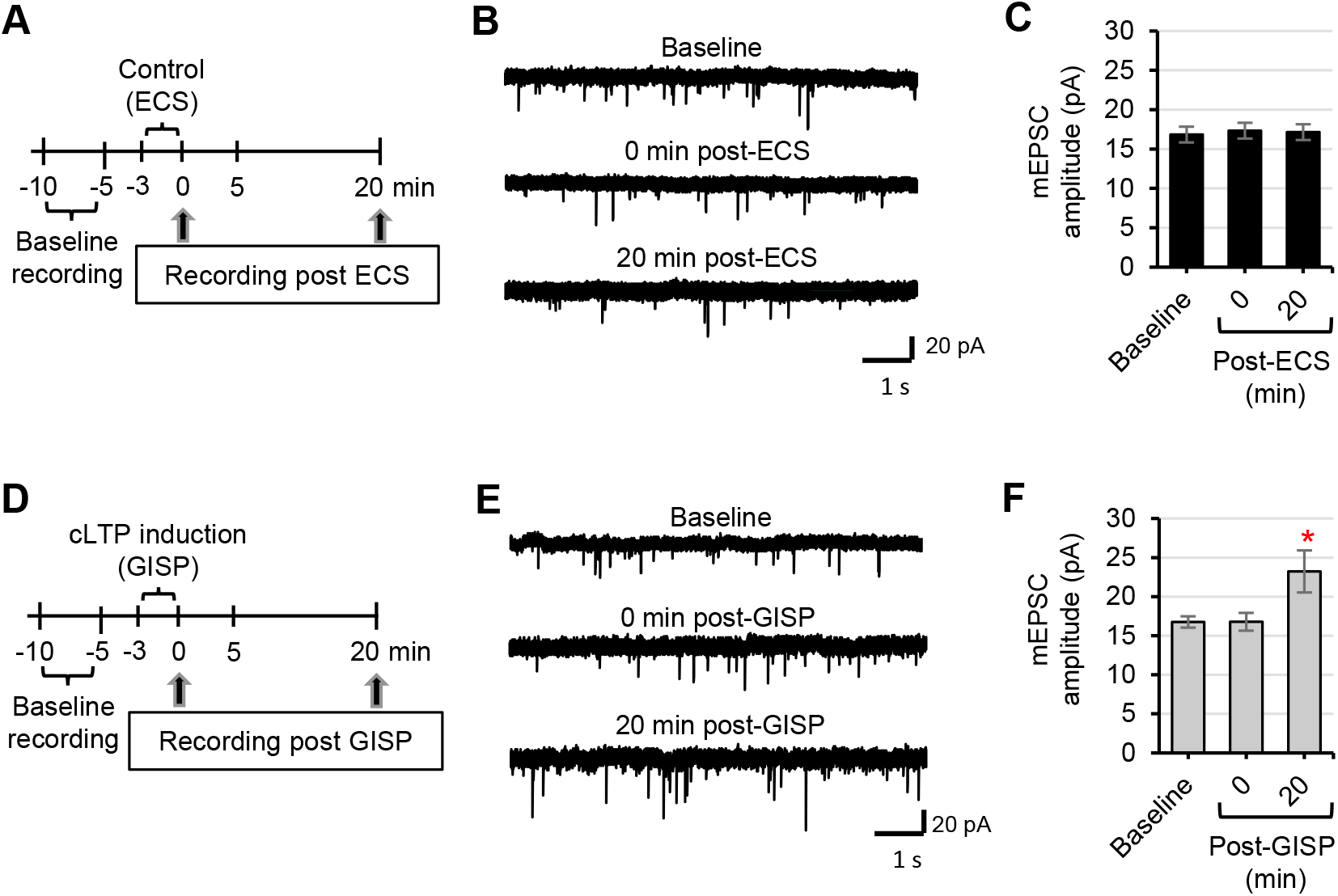
Electrophysiological characterization of cLTP expression in primary hippocampal cultured neurons. **(A, D)** Schematic of cLTP induction protocol used in the mEPSC recording. Whole-cell patch clamp is made with cultured hippocampal neurons (DIV 15) and mEPSCs are recorded for 5 min as baseline. Neurons were incubated for 3 min in artificial cerebral spinal fluid as extracellular control solution (ECS)**(A-C)** or glycine-induced synaptic potentiation (GISP) buffer containing glycine (200 μM) in ECS without Mg^2+^ to trigger cLTP **(D-F)**. After GISP buffer, neurons were returned to the ECS for 0 and 20 min. **(B, E)** Representative mEPSC traces. **(C, F)** Quantification of the mEPSC amplitude at baseline, 0, and 20 min after ECS or GISP treatment. GISP treatment potentiates the mEPSC in primary hippocampal neurons. Statistical significance is determined using ANOVA. n = 10 neurons per treatment (*p < 0.05).

**Figure S2:**
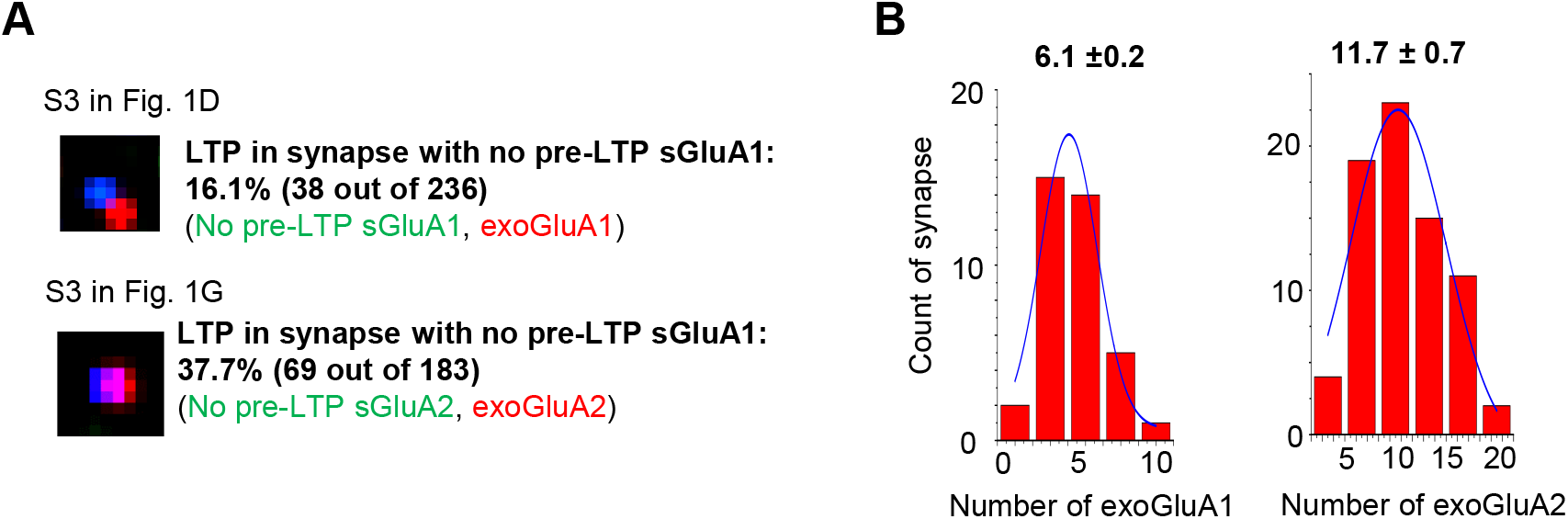
Quantification of synapses that had no surface AMPAR subunits before cLTP induction but gained them after cLTP induction. **(A)** Zoomed images of Synapse S3 in Fig. 1D and Synapse S3 from Fig. 1G. These synaspes show a PSD marker Homer1c-mGeos (blue) and newly exocytosed GluA1 and GluA2 (exoGluA1 and exoGluA2) induced by cLTP induction (red), but not surface GluA1 and GluA2 before cLTP induction (pre-LTP sGluA1 or sGluA2) (green). **(B) Quantification of synapses that show LTP but no pre-existing surface AMPAR subunit prior to LTP, in relation to the number of exocytosed subunit**. A total of 236 synapses for GluA1 was measured. A total of 183 synapses for GluA2 was measured.

**Figure S3.**
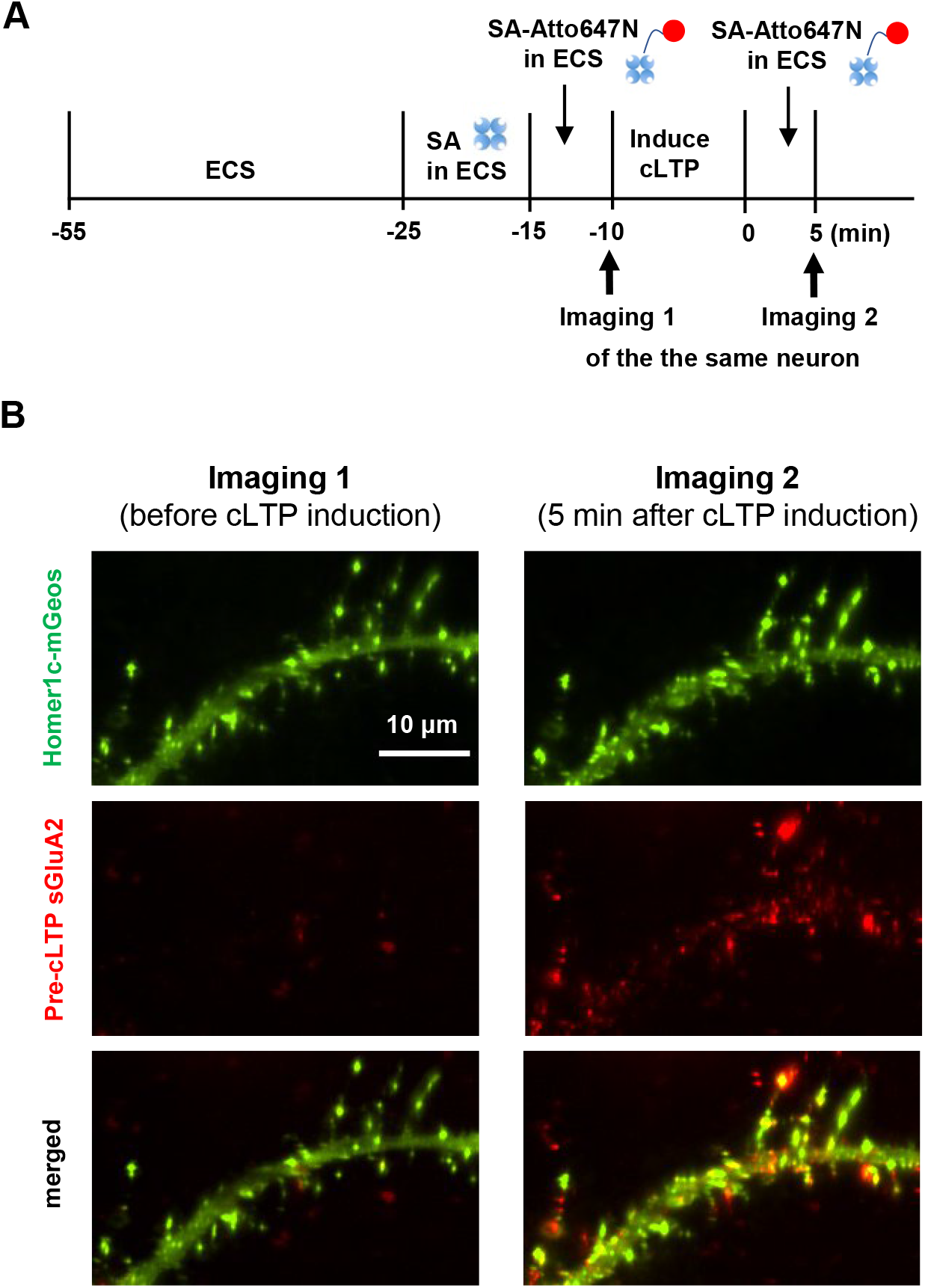
Visualization of AMPAR insertion upon cLTP induction on single synapses basis. **(A)** Experimental sequence of labeling biotinylated surface AMPAR subunit using Atto647N-conjugated SA (SA-Atto647N) before and after cLTP induction protocol for 10 min. Neurons were transfected with Homer1c-mGeos, GluA2 containing AP tag which were biotinylated by cotransfected BirA. Neurons were incubated with extracellular control solution (ECS) for 30 min and then with 10 nM unconjugated SA in ECS to saturate the labeling of the existing surface biotinylated GluA2 before cLTP induction. To check the efficiency of the labeling with unconjugated SA, neurons were incubated with 0.6 nM SA-Atto647N in ECS for 5 min, then immediately imaged. To visualize the extent of AMPAR insertion after cLTP induction, neurons were incubated with 200 μM glycine in GISP solution without Mg2+ for 10 min to induce cLTP, followed by application of 0.6 nM SA-Atto647N for 5 min and imaging. **(B)** Representative images of surface biotintylated GluA2 subunits labeled by SA-Atto647N after applying unconjugated SA and after applying cLTP induction protocol in the same neuron. Minimal SA-Atto647N labeling at imaging 1 indicates that the labeling of pre-existing surface biotinylated GluA2 by 10 nM SA reached saturation. Strong SA-Atto647N labeling in the same neuron after cLTP induction in imaging 2 indicates cLTP induction protocol induced AMPAR insertion within 5 min.

**Figure S4.**
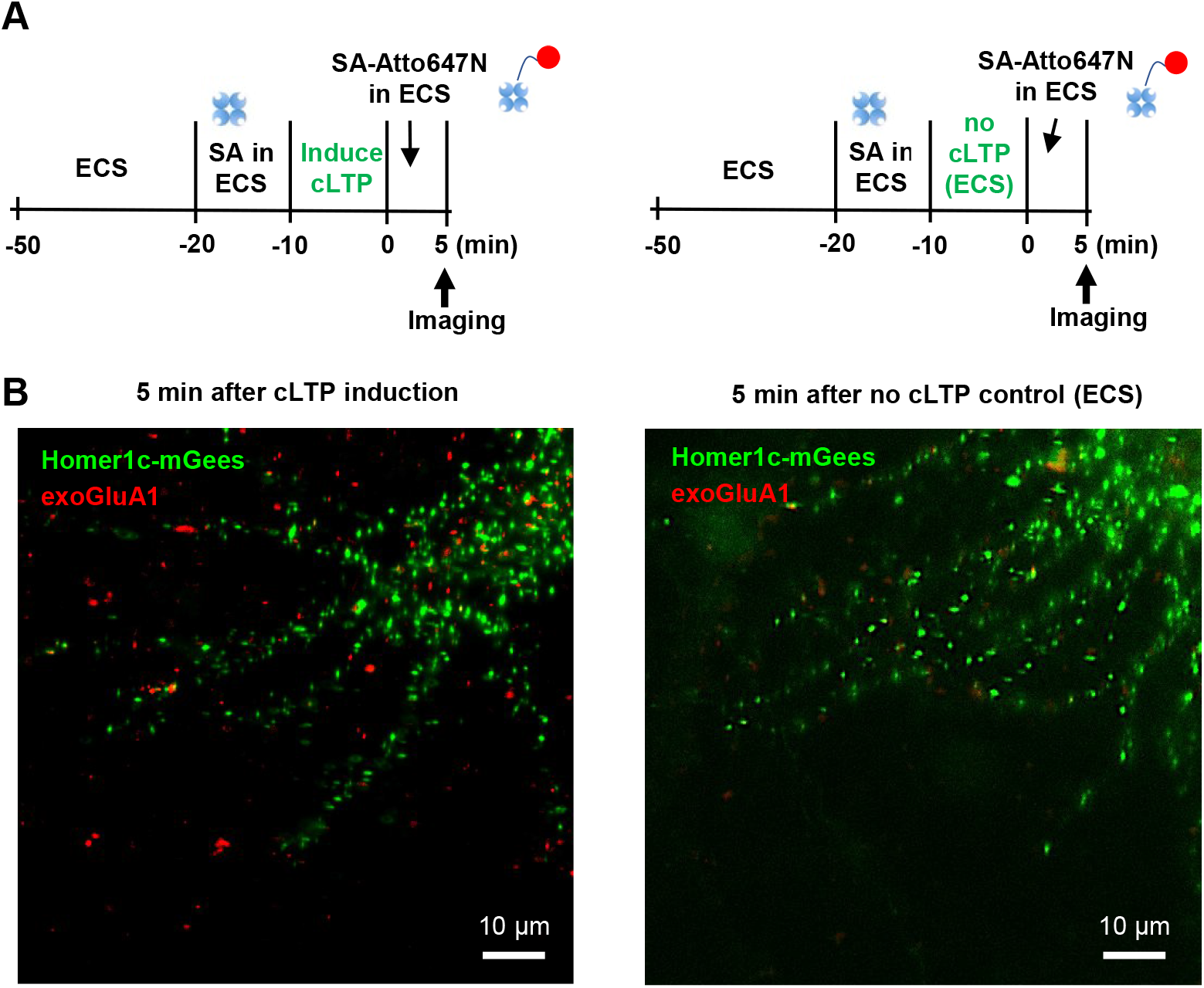
AMPAR insertion upon cLTP induction is the major source of increase in Atto647N signal. **(A)** Experimental sequence of labeling biotinylated surface AMPAR subunit using Atto647N-conjugated SA (SA-Atto647N) after cLTP induction protocol or control ECS incubation for 10 min. Homer1c-mGeos was co-transfected in neurons as a PSD marker. **(B)** Representative images of newly inserted GluA1 subunits labeled by SA-Atto647N after cLTP induction protocol or control ECS incubation. This cLTP induction protocol induced exocytosis of transfected GluA’s within 5 min. However, <5% of synapses had exocytosed GluA2 in neurons treated with control ECS solution (9 out of 200 synapses counted), demonstrating a significant difference between ECS and GISP treatment.

**Figure S5.**
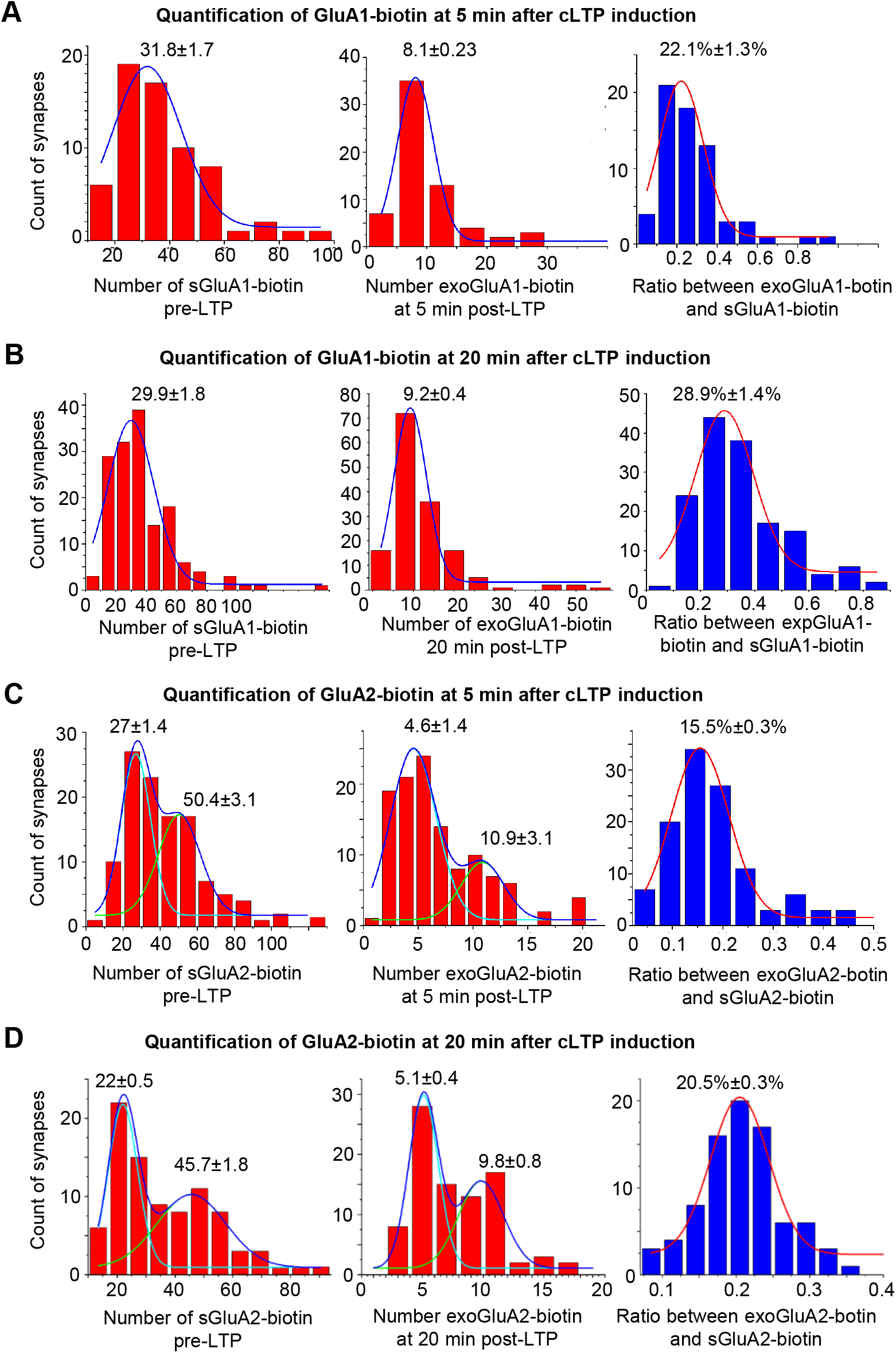
On transfected neurons, newly inserted GluA1 and GluA2 differently contribute to the synaptic AMPAR level during the cLTP maintenance phase. **(A-B)** The quantification of the number of surface-GluA1 before cLTP induction (pre-LTP sGluA1) and newly-inserted GluA1 (exoGluA1) after cLTP induction at 5 min **(A)** and 20 min **(B)** after cLTP induction. **(C-D)** The quantification of the number of surface-GluA2 before cLTP induction (pre-LTP sGluA2) and newly-inserted GluA2 (exoGluA2) after cLTP induction at 5 min **(A)** and 20 min **(B)** after cLTP induction. A total of 65 synapses for GluA1 was measured from 3 neurons in 2 independent experiments at 5 min post cLTP induction. A total of 151 synapses for GluA1 was measured from 3 neurons in 2 independent experiments at 20 min post cLTP induction. A total of 116 synapses for GluA2 was measured from 3 neurons in 2 independent experiments at 5 min post cLTP induction. A total of 88 synapses for GluA2 was measured from 3 neurons in 2 independent experiments.

**Figure. S8.**
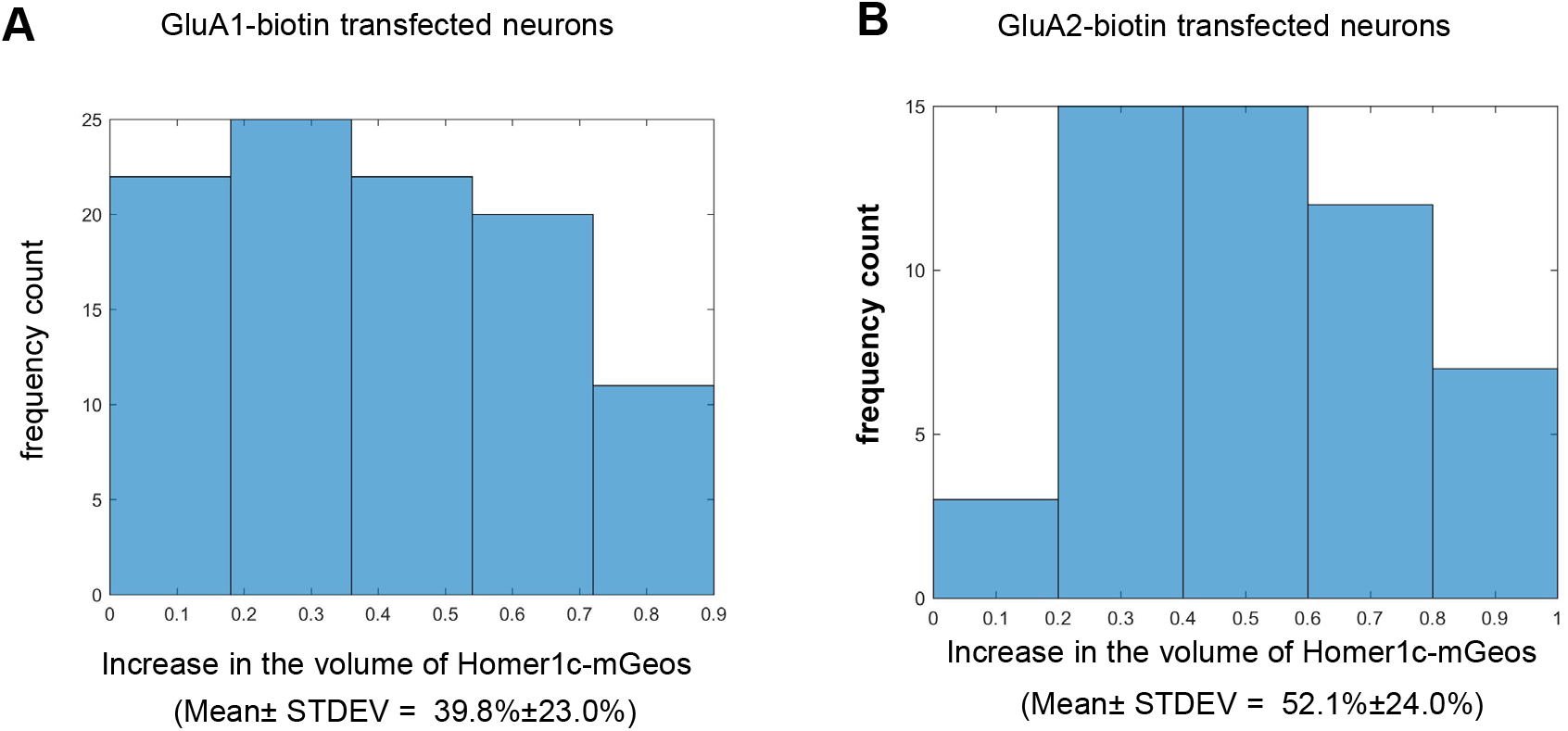
The induction of cLTP Increases the volume of Homer1c-mGeos fluorescence intensity. **(A)** Frequency count histogram on *synapses* in relation to the increase in the volume of Homer1c-mGeos in GluA1-biotin transfected neurons in Fig. 3. The average increase in volume of Homer1c-mGeos is 39.8 ± 23.0 % (Mean ± STDEV) computed from 100 synapses in 3 transfected neurons from 3 independent cultures. **(B)** Frequency count histogram on *synapses* in relation to the increase in the volume of Homer1c-mGeos in GluA2-biotin transfected neurons in Fig. 3. The average increase in volume of Homer1c-mGeos is 52.1 ± 24.0 % (Mean ± STDEV) computed from 52 synapses in 2 transfected neurons from 2 independent cultures.

**Figure S7.**
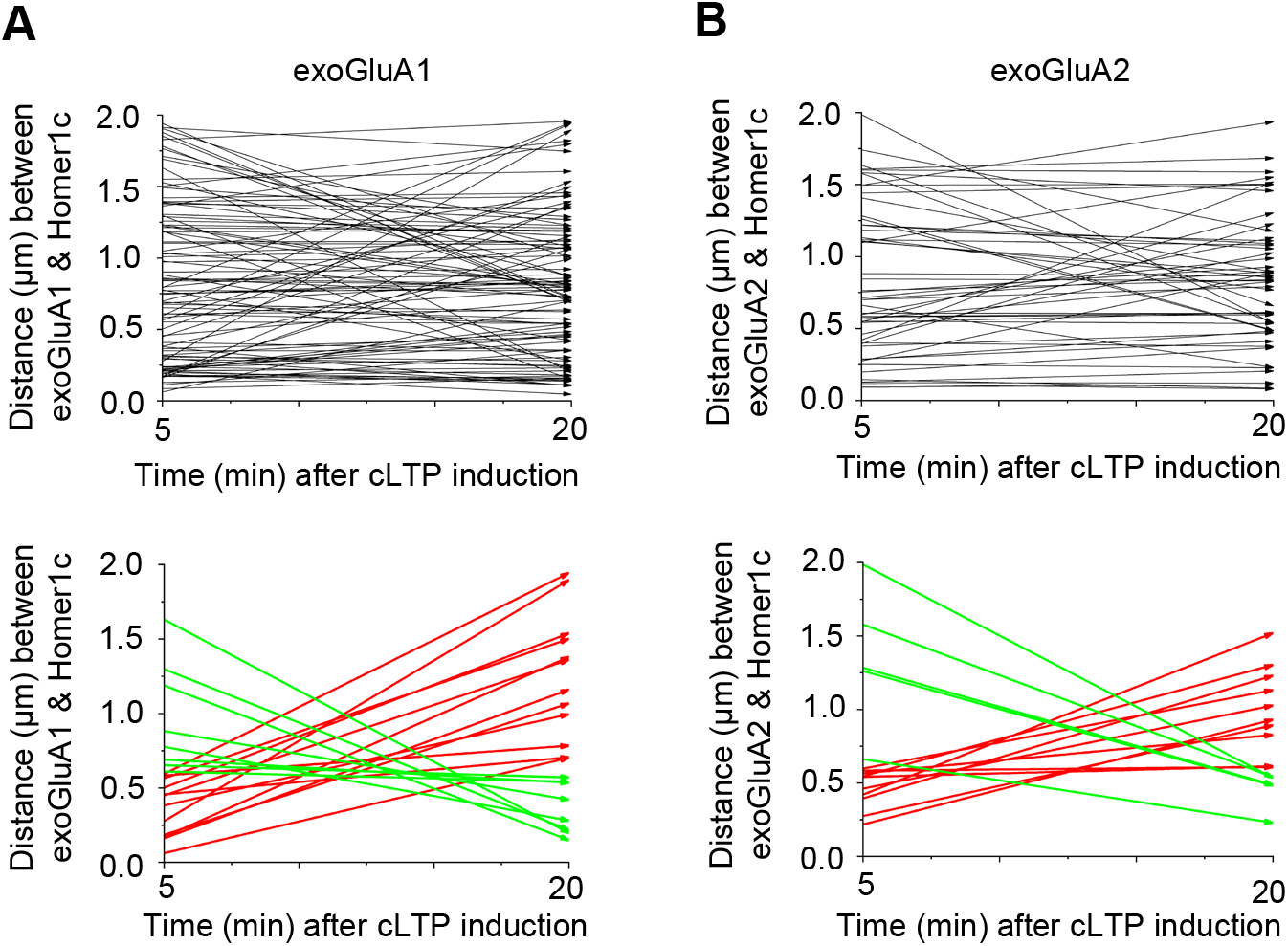
The exchange of exocytosed AMPARs between synaptic and extrasynaptic regions does not significantly change during 5-20 min post cLTP induction when the synaptic region was defined as being 0.6 µm from the center of Homer1c. This figure is supplemental to Fig. 3. The synaptic region was defined as being 0∼0.6 µm from the center of Homer1c, and the distance between AMPAR subunit and the center of Homer1c was recalculated. The end of an arrow stands for the moving direction of the subunit. Distance between the center of Homer1c and GluA1 **(A)** and GluA2 **(B)** at 5 min and 20 min after cLTP induction. The top graphs show all traces (black). The bottom graphs display the traces extracted from the top graphs, which show the subunits moving from synaptic to extrasynaptic region (red), or those from extrasynaptic to synaptic region (green). **(A)** Out of 100 GluA1 from transfected neurons there are 12 moving from synaptic to extrasynaptic region, and 8 moving from extrasynaptic to synaptic region. **(B)** Out of 52 GluA2 from transfected neurons, there are 9 GluA2 moving from synaptic to extrasynaptic region, and 5 moving from extrasynaptic to synaptic region.

**Figure S8.**
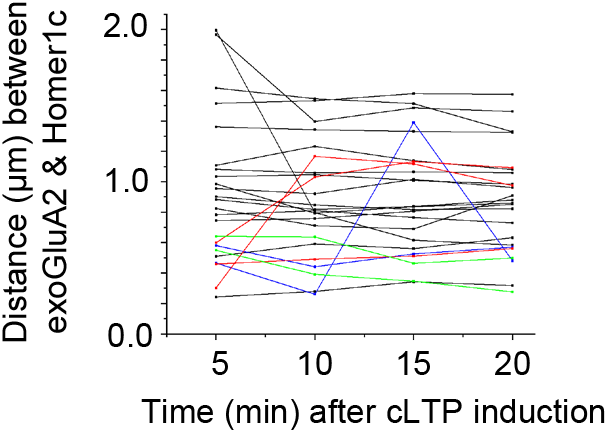
The exchange of exocytosed AMPARs between synaptic and extrasynaptic regions does not significantly change during 5-20 min post cLTP induction when they are labeled with mSA-sQDs. To validate our results with SA-Atto647N in Fig. 3, we repeated the experiments outlined in Fig. 3B using small QDs conjugated with monomeric SA (mSA-sQDs), which allowed imaging every 5 min from 5-20 min after cLTP induction due to their photostability. The synaptic region was defined as being 0∼0.5 µm from the center of Homer1c, whereas the extrasynaptic region was defined as being 0.5-2 µm from the center of Homer1c. The distance between the center of Homer1c and exocytosed GluA2 was measured at 5, 10, 15, and 20 min after cLTP induction. Out of the 24 GluA2 traces, 17 did not cross the 0.5 µm mark from the center of Homer1c (black). Out of 7 that crossed the 0.5 µm mark, 2 went from extrasynaptic to synaptic (green) and 3 went from synaptic to extrasynaptic region (red). There were also 2 that moved to both synaptic and extrasynaptic regions (blue).

## Supplementary Tables

**Table S1.** The % increase of transfected GluA1 and GluA2 on neuronal plasma membrane at 5 min and 10 min after cLTP induction compared to their basal surface expression before cLTP induction.

|  | GluA1 | GluA2 |
| --- | --- | --- |
| # of sGluA | 32 | 27 or 50 |
| increase in GluA @ 5 min<br>after cLTP compared to<br>basal level<br>(synaptic+extrasynaptic) | 22.2% (of this 22.2%, 34%<br>synaptic, 66% extrasynaptic) | 15.5% (of this 15.5%, 32% synaptic,<br>68% extrasynaptic) |
| increase in GluA @ 20 min<br>after cLTP compared to<br>basal level<br>(synaptic+extrasynaptic) | 28.9% (of this 28.9%, 48%<br>synaptic, 52% extrasynaptic) | 20.5% (of this 20.5%, 50% synaptic,<br>50% extrasynaptic) |

**Table S2.** Changes in the number of transfected GluA1 and GluA2 on neuronal plasma membrane at 5 min and 10 min after cLTP induction compared to their basal surface expression before cLTP induction.

|  | GluA1 |  | GluA2 |  | GluA2 |  |
| --- | --- | --- | --- | --- | --- | --- |
|  | 5 min after cLTP | 20 min after cLTP | 5 min after cLTP | 20 min after cLTP | 5 min after cLTP | 20 min after cLTP |
| increase in synaptic GluA compared to basal level | 2.4 | 4.5 | 2.5 | 5 | 1.3 | 2.75 |
| increase in extrasynaptic GluA compared to basal level | 4.6 | 4.8 | 5.3 | 5 | 2.9 | 2.75 |
| Total increase | 7<br>(32*22.2%) | 9.3<br>(32*28.9%) | 7.8<br>(50*15.5%) | 10<br>(50*20%) | 4.2<br>(27*15.5%) | 5.5<br>(27*20.5%) |

**Table S3.** Summary of the diffusion constants of GluA1 and GluA2 at synaptic and extrasynaptic region at 5 min and 20 min after cLTP induction. The synaptic region was defined as being 0∼0.5 µm from the center of Homer1c, whereas the extrasynaptic region was defined as being 0.5-2 µm from the center of Homer1c.

| Diffusion Constant at 5 min post cLTP induction (1000*$\mu\text{m}^2/\text{s}$) | | | |
| --- | --- | --- | --- |
|  | synaptic | extrasynaptic | Fold Difference |
| GluA1 | 8.4±0.46 | 11.2±0.67 | extrasynaptic is 1.33 times of synaptic |
| GluA2 | 6.1±0.12 | 12.7±1 | extrasynaptic is 2.08 times of synaptic |
| Diffusion Constant at 5 min post cLTP induction (1000*$\mu\text{m}^2/\text{s}$) | | | |
|  | synaptic | extrasynaptic | Fold Difference |
| GluA1 | 13±1 | 22.4±0.64 | extrasynaptic is 1.72 times of synaptic |
| GluA2 | 8.1±0.17 | 19±2.3 | extrasynaptic is 2.35 times of synaptic |

